# BioMetAll v2.0: Introducing Scores, Metal Discrimination, and Side-Chain Descriptors for Predicting Metal-Binding Sites in Proteins

**DOI:** 10.64898/2026.07.09.737562

**Authors:** José-Emilio Sánchez-Aparicio, Raúl Fernández Díaz, Raúl Peña Losada, Weifeng Gao, Mercè Alemany Chavarria, Jean-Didier Maréchal

## Abstract

Predicting the location of metal-binding sites in proteins is crucial for fundamental biological questions and biotechnological applications. Over the past decade, the rise in metal-bound protein structures in the Protein Data Bank, combined with advanced statistical models such as deep learning, has accelerated the development of metal-binding site prediction tools. Several approaches are now available, offering high-quality benchmarks and predictive performance. Our initial development in this area is BioMetAll, whose first version was based on backbone pre-organization. Here, we introduce its second version, featuring two major updates: 1) metal-specific scoring functions and 2) prediction using backbone geometry alone or in combination with first coordination sphere descriptors. Apart from demonstrating metal sensitivity and yielding better benchmarking results, this new version allows the assessment of the influence of considering the metal’s first coordination sphere versus backbone pre-organization on how metallic species bind to proteins.

## Introduction

More than 30% of biological systems recruit metals for structural or catalytic purposes.^1^ The diversity of functions exhibited by metalloproteins constantly inspires chemists, and research endeavors into bioinorganic interactions abound. Their aim includes deciphering the intricate molecular machinery of nature and engineering novel biometallic species. In this field, predicting how metals interact with biological scaffolds at the molecular level is particularly relevant.^2^ Several computational strategies exist with this objective. All pretend to assess, from structure and/or sequence, the location of the metal, which amino acids bind to the metal, and the geometry of the resulting coordination sphere.^3,4^

Notwithstanding recent efforts, the available pool of predictors remains relatively limited, with substantial room for further advancement. Thus far, most software relied on matching templates to well-established binding sites or metal-binding sequences,^5–10^, with recent ones based on machine learning and genetic algorithms.^11–17^ However, in these software solutions, the focus is mainly on reproducing metal-binding sites that have already been identified within proteins. This approach offers limited flexibility for generating entirely new structural configurations or looking for sites that are not in ideal geometries (e.g., transient or incipient ones).

In 2021, we proposed an alternative strategy embedded in a new software package called BioMetAll.^18^ The rationale of the approach is based on the preorganization hypothesis: the backbone of each amino acid involved in metal coordination accounts for compatible geometries for metal binding. Each amino acid, therefore, has a specific geometric subspace of the backbone to define for the side chain to organize itself toward metal binding. To construct BioMetAll, relatively simple geometric descriptors were used as markers of backbone preorganization (e.g., distances from the metal to the alpha carbon of the coordinating amino acids), and the library was built from statistics on more than 100000 reported crystallographic binding sites. BioMetAll uses a workflow that begins by embedding the protein in a grid of probes. Those probes that meet all the geometric criteria for a certain number of amino acids are selected as potential for metal coordination. Finally, the program generates the metal-binding sites by merging probes that share the same amino acid coordination environment (i.e., the same set of amino acids that can form coordination bonds with the metal).

The original BioMetAll algorithm was carefully designed to support *de novo* design efforts. This led to a primary focus on identifying metal-binding regions rather than precise coordination spheres. Within this paradigm, BioMetAll 1.0 had two main limitations. The first one comes from the Boolean evaluation of the probes: a probe is assigned a value of 1 if it meets the geometric requirements, and 0 otherwise. Although this approach has been successful in identifying both stable and even transient metal-binding sites in proteins, it may, in some cases, yield false-positive locations.^11^ The second issue is that no differentiation between metals was available. This limitation could be partially addressed by manually configuring the search to identify amino acids that are preferential for coordinating a given metal. However, hindsight revealed that it would be more practical to automatically search for metal-specific binding sites. Additionally, regarding other predictors, BioMetAll does not allow consideration of pure first-coordination-sphere participation; a refinement that could be necessary in some cases.

To address these issues, here we introduce a new version of BioMetAll (installation instructions and open-source code are available at https://github.com/insilichem/biometall2). This BioMetAll 2.0 has major advances that include: i) a new scoring function to reduce false positives and better discriminate the most stable sites, ii) a new descriptor –coordination bond distance– accounting for the side-chain preorganization, and iii) metal specificity that includes transition metals, alkaline, and alkaline earth. To demonstrate the method’s improvement, we benchmarked the new version and observed substantial gains over the original BioMetAll.

## BioMetAll improvements

### The new scoring function allows for better discrimination of metal-binding sites

At the core of the new BioMetAll’s capabilities is the incorporation of a scoring function to evaluate and rank the metal-binding sites predicted by the program. The hypothesis is that the new scoring function will allow the most stable sites (e.g., those present in crystallographic structures) to be better positioned in the ranking than under the previous criterion, which relied only on the number of probes each site has.

To build the scoring function, we have relied on two assumptions: i) the proportion of coordination geometries available in current experimental databases is a fair representation of their prevalence in nature; and ii) there is a direct correlation between the prevalence of a certain coordination geometry in nature and how favorable it is. To differentiate between metals, we propose a similar hypothesis: the affinity of a specific metal for a residue correlates with the presence of this metal-residue interaction in the databases. The underlying idea is therefore to award higher scores to those coordination geometries that appear more frequently in the database and to punish those that are rarer. This goal has been achieved through a series of statistical analyses on the same dataset used in the original version of BioMetAll.^18– 20^ In the following paragraphs, we detail the algorithm’s scoring process.

Given a metal type (Zn, Fe, Mn, Cu, Cd, Ni, Co, Hg, Ca, Mg, Na, K, as well as a “Transition” atom type), we first measure the probability that a probe will be bound to a certain amino acid. For now, to model this probability, we continue using the geometric features from the previous version of the program. There are two cases: coordination with a donor atom on the side chain and coordination with the backbone oxygen atom as the donor. For side-chain coordination, three descriptors are used: i) distance from the probe to the alpha carbon of the amino acid, ii) distance from the probe to the beta carbon of the amino acid, and iii) angle formed between the probe, the alpha carbon, and the beta carbon. For the backbone-oxygen coordination, two descriptors are used: i) distance from the probe to the backbone-oxygen atom, and ii) angle formed between the probe, the backbone-oxygen atom, and the backbone-carbon atom. For each triad descriptor/metal type/residue type, we constructed its probability density function and adjusted it to be bimodal to account for the two “peaks” observed in some cases. The details of entire set of bimodal functions are provided in the repository annexed (directory 01_bimodal_functions). To obtain the score of a probe-amino acid pair, we calculate the mean of the bimodal functions applied to the descriptor values. An example of Zn/Glu side-chain coordination is shown in Figure 1.

**Figure 1.**
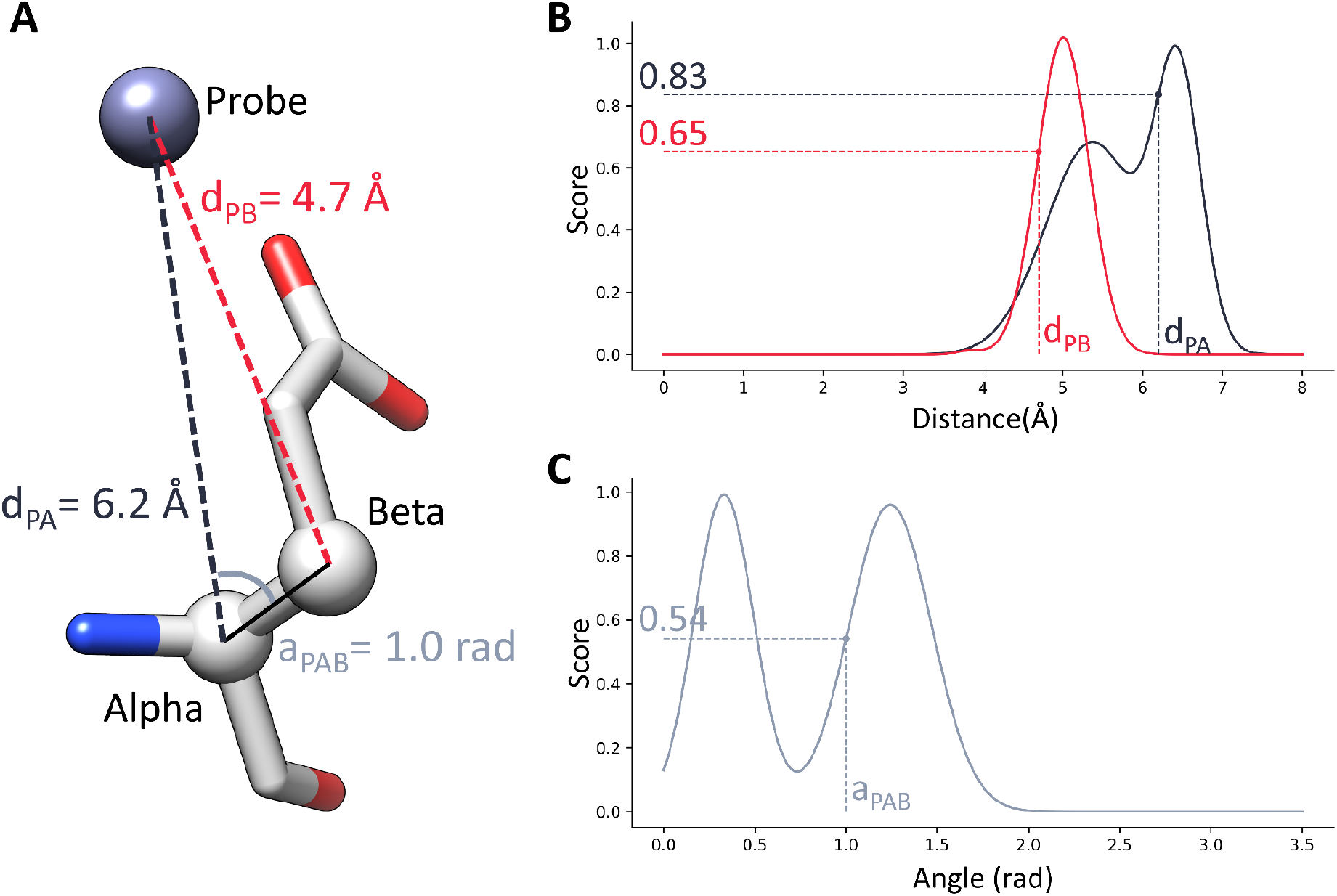
Example of score calculation for a probe in the Zn/Glu case. (**A**) Visual representation of the probe-amino acid geometry. Values of the three descriptors (distance probe-alpha carbon, distance probe-beta carbon, and angle probe-alpha carbon-beta carbon) are highlighted in dark gray, red, and light gray, respectively. (**B**) The probability density function for the distance between the probe alpha carbon and the probe is shown in dark gray. The partial score for the example is 0.83. The probability density function for the distance between the probe and beta-carbon is shown in red. The partial score for the example is 0.65. (**C**) The probability density function for the angle probe (alpha-carbon-beta-carbon) is shown in light gray. The partial score for the example is 0.54. The final score is calculated by averaging the three partial scores (result = 0.67). Further weighting by amino acid prevalence and normalization is performed.

After calculating scores for all probe-amino acid pairs, they were weighted by the frequency with which the metal is bound to that residue in the database. They were then normalized within the range [0, 1], considering the maximum possible score for that metal (provided in the repository annexed in directory 02_score_normalization). Probe-amino acid pairs with non-negligible scores are chosen as potential components of a metal-binding site.

The final steps of the algorithm aim to build the list of predicted metal-binding sites: i) check those probes that present coordination with a set of amino acids compatible with the user’s requirements (e.g., have a minimum number of amino acids involved or match a specific motif); ii) group all the probes that belong to the same coordinating environment (i.e., metal-binding site); iii) assign a score to each metal-binding site, which will be the maximum probe score among all those assembling the site; and iv) output a list of the predicted metal-binding sites ordered by score.

To assess whether incorporating the score improves the predictions of the original BioMetAll, we used the same motif-search benchmark of 53 protein structures. The task was to detect sites where the metal could be bound by a two-histidine one carboxylate motif (FTM). In this benchmark, we instructed BioMetAll 2.0 to search for all possible FTMs for a “Transition” metal to avoid metal specificity and allow a fair comparison with the previous version of the program. Taking the most demanding metric, which is to find the exact crystallographic motif at the first position of the predicted sites, results show an improvement in the success rate from 40% in BioMetAll 1.0 to 62% in BioMetAll 2.0. This result indicates that incorporating the score allows discrimination of the most stable sites (i.e., crystallographic) from the more transient or incipient ones. Noticeably, in BioMetAll 2.0, the distance from the predicted position of the metal (highest-scored probe) to the crystallographic one is, on average, 1.0 ± 1.0 Å, whereas in BioMetAll 1.0, the most centered probe did not allow for accurate prediction of the position of the metal (distance 2.5 ± 0.8 Å). Altogether, the results of this benchmark support the hypothesis that a large part of the metal location is explained by the protein’s backbone preorganization. However, a fraction of cases still show that the backbone descriptors were insufficient to discern the most stable sites.

### First-sphere preorganization is useful to identify stable metal-binding sites

To improve the prediction of the most stable metal-binding sites, we wondered whether introducing a new descriptor that accounts for atoms in the first coordination sphere could be useful. When looking at state-of-the-art predictors, all use the coordinates of side-chain atoms to make predictions, either to build templates used by the algorithm to generate predicted binding sites or to train the neural network responsible for the prediction.^11-17^ Although the introduction of a “first-sphere” descriptor in Biometall implies a further step beyond the backbone preorganization hypothesis, it seems reasonable that it could help to refine the results.

Keeping in mind the simplicity of all the other descriptors used in BioMetAll, we thought that measuring the “coordination distance” (i.e., distance from the probe to the side-chain donor atom) would be the best way to proceed. Therefore, the complete landscape of descriptors when the donor atom belongs to the amino acid side-chain (Figure 2) includes the three features related to the alpha and beta carbons (denominated “AB” in the program parameters) and the new descriptor (denominated “FS”) to account for the first sphere of metal coordination.

**Figure 2.**
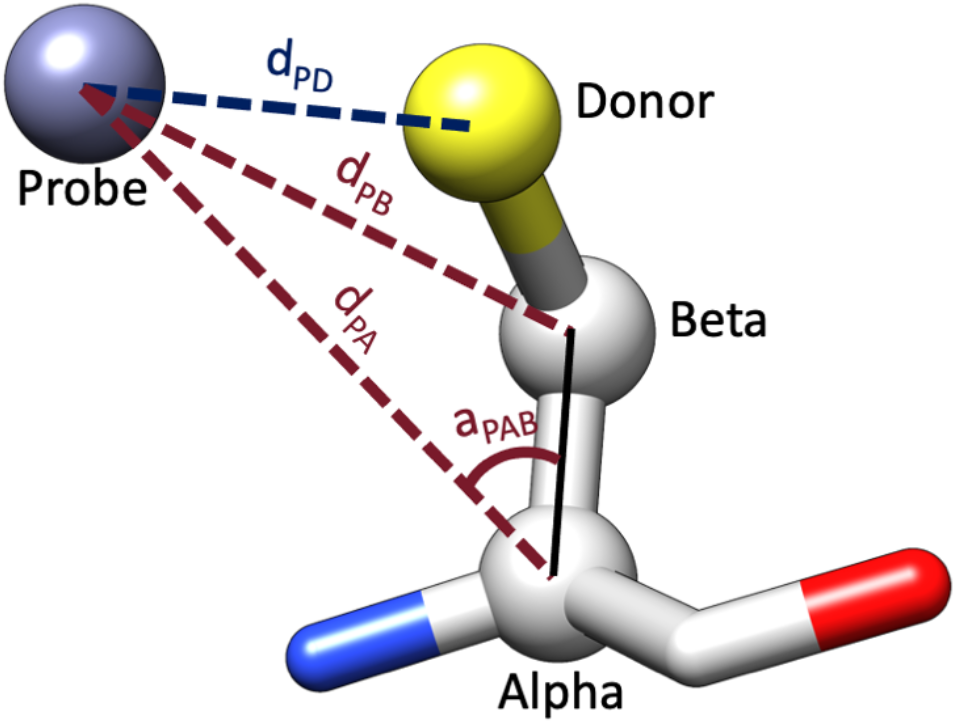
Descriptors to evaluate coordinations with a side-chain donor atom. Descriptors related to alpha and beta carbons are shown in dark red: distance from the probe to the alpha carbon of the amino acid (d_PA_), distance from the probe to the beta carbon of the amino acid (d_PB_), and angle formed between the probe, the alpha carbon and the beta carbon (a_PAB_). The new descriptor accounting for the distance between the probe and the donor atom (d_PD_) is shown in dark blue.

To implement the score for the new “FS” descriptor, we followed the same approach as for the “AB” descriptors. First, we measured the coordination distance for all metal-donor bonds in the database. Then, for each pair of metal type/residue type, we constructed the probability density function for the coordination distance descriptor and adjusted it to be bimodal (details are provided in the directory *01_bimodal_functions* of the repository annexed). The “FS” score is therefore the bimodal function applied to the probe-donor distance, followed by residue prevalence weighting and normalization. Finally, to obtain the score of a probe-amino acid pair when using both backbone and side-chain related features, we calculate the mean of “AB” and “FS” scores.

We repeated the motif search benchmark, instructing BioMetAll 2.0 to find FTMs for a “Transition” metal in the same set of 53 structures. In this case, the program considered both the “AB” and “FS” descriptors when scoring the metal-binding sites. Again, to evaluate the importance of the “FS” descriptor, we consider the metric of finding the exact crystallographic motif at the first position of the predicted sites. Results show an improvement in success rate from 62% with only “AB” descriptors to 96% with both “AB” and “FS” (details are reported in the directory *03_benchmark_vs_BioMetAll1*.*0* of the repository annexed). Incorporating coordination distance into scoring greatly reduced the number of results (from an average of 9 ± 9 sites in the “AB” case to 2 ± 1 sites in the “AB” + “FS” case). The average distance between the predicted and crystallographic positions of the metal (highest-scored probe) is also substantially improved, from 1.0 ± 1.0 Å to 0.7 ± 0.7 Å.Taken together, we conclude that incorporating first-sphere preorganization into the scoring enabled nearly complete identification of crystallographic FTMs, including prediction of metal locations.

PDB code 3n9d^21^ was the only crystallographic FTM not predicted using “AB” + “FS” descriptors. When only the “AB” descriptors were used, three metal-binding sites were identified (Figure 3A). The best-scored corresponds to the FTM crystallographic triad (His42, His48, Asp50) and has a relatively low score of 0.274. A second site with a score of 0.243 is identified in an area more exposed to the solvent (Asp117, His127, His142), and a third site (His42, Asp50, His84) with a very low score (0.115) near the crystallographic motif. Visual inspection of the results confirms that the first and second sites exhibit viable backbone geometries for coordination, whereas the third site appears much more forced, as reflected in the score.

**Figure 3.**
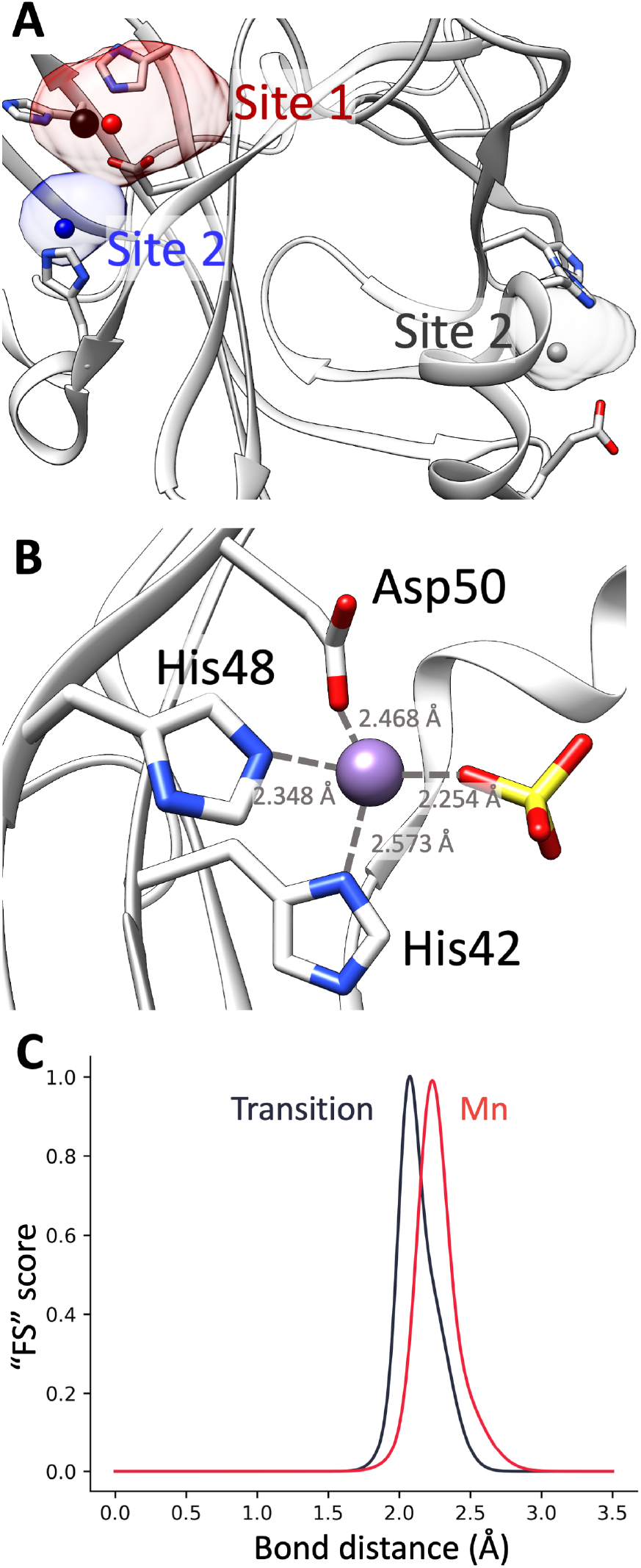
(**A**) Predicted metal-binding sites using “AB” descriptors for the case of a transition metal in PDB 3n9d. Sites 1, 2 and 3 are depicted in red, blue and gray, respectively. Best-scored probes are shown as spheres. The crystallographic position of the metal is shown as a black sphere. (**B**) Crystallographic coordination geometry detail. (**C**) Evaluation function for the “FS” descriptor in the case of histidine coordination. In dark gray is shown the function for a transition metal, and in red for a manganese ion.

When the “FS” descriptor is added, no viable metal-binding sites are predicted. If we closely inspect the crystallographic coordination geometry (Figure 3B), we observe that coordination distances between the metal and the three amino acids involved are 2.57 Å(His42), 2.35 Å (His48), and 2.47 Å(Asp50). We also observe the presence of an exogenous ligand (sulfate). While the scoring evaluation of coordination distances shows acceptable results for His48 and Asp50, it is not the case for His42, which has a very low score (Table 1). Presumably, BioMetAll did not find any probe with a convenient distance to His42 and therefore discarded this option.

**Table 1.**
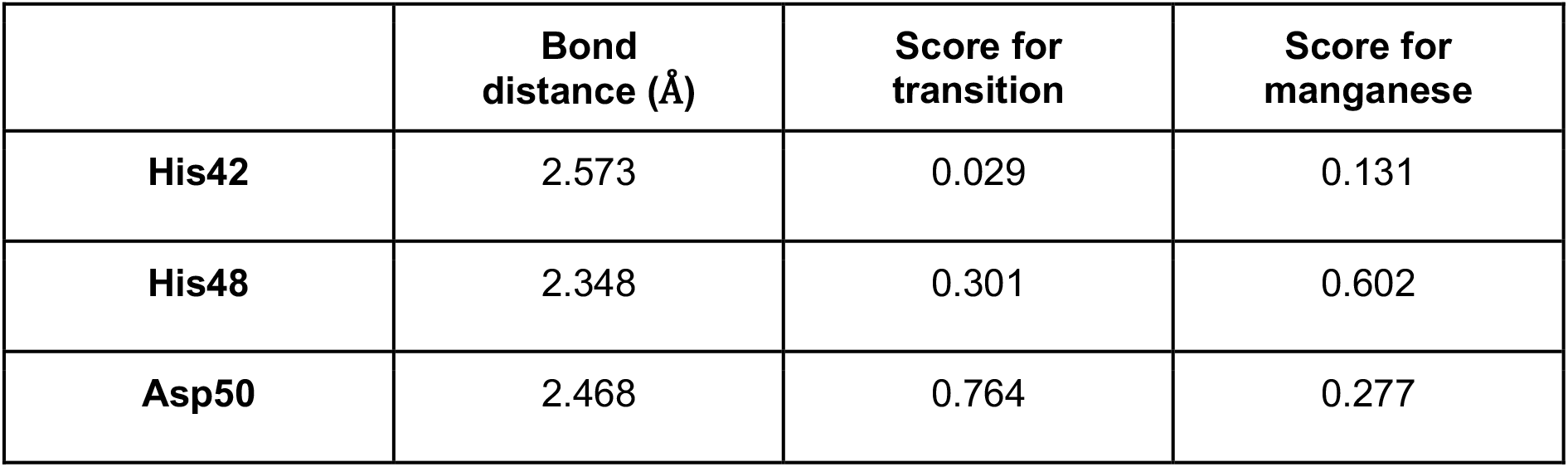
Individual “FS” scores of each amino acid in 3n9d FTM, considering the crystallographic metal position and two sets of parameters: for a transition metal and for a manganese ion.

|  | **Bond distance (Å)** | **Score for transition** | **Score for manganese** |
| --- | --- | --- | --- |
| **His42** | 2.573 | 0.029 | 0.131 |
| **His48** | 2.348 | 0.301 | 0.602 |
| **Asp50** | 2.468 | 0.764 | 0.277 |

It should be noted that the metal involved in the coordination of 3n9d FTM is a manganese ion, which represents a low percentage (7%) of the bonds included in the statistical analysis for the “Transition” case.^18^ We aimed to ascertain if the absence of results was due to the presence of the sulfate (which could distort the coordination geometry) or to the specificities of the metal type. To do so, we repeated the calculation using manganese parameters rather than the ones for transition metals. In this case, the crystallographic site was identified as the only solution, with a score of 0.574 and a distance of 0.6 Å between the best-scoring probe and the crystallographic metal. Analysis of the individual “FS” scores for the crystallographic metal (Table 1) shows that all three amino acids receive a non-negligible score, suggesting that the main reason for the absence of results in the “Transition” exploration was the particularities of the probability function of the His-Mn bond distance with respect to that of His-Generic (Figure 3C).

The incorporation of the scoring function and the new coordination-distance descriptor enables much better prediction and discrimination of stable metal-binding sites. Analysis of case 3n9d reveals some characteristics of BioMetAll 2.0 beginning to emerge. First, the user can choose among different sets of descriptors, each offering a different view of the protein preorganization required to form a metal-binding site. Second, the ability to select from a wide set of metal types and generic offers greater customization in terms of metal specificity. Given the program’s greater flexibility and the relatively low computational cost of the calculations (an average of 11 s per run on the FTM benchmark on a desktop personal computer), we encourage users to explore different parameter settings when studying a system. In the following sections, we will go deeper into this sense.

### Tolerance parameterization

A consequence of introducing the score is the difficulty in establishing a threshold to distinguish between “good” and “bad” solutions (i.e., candidates for a stable site versus more transient ones). Despite the weighting and normalization applied at the end of the scoring process, the score may remain insufficiently informative for the user to make a decision. To overcome this obstacle, we introduced the concept of “tolerance,” which is closely related to the traditional terms of precision and recall, widely used in machine learning to evaluate and parameterize algorithms.

We parametrized three predefined tolerance levels for each metal: low, medium, and max. The user could bypass these cutoffs by defining a custom score (accessible via the –score *score_value*). By default, the program employs a medium tolerance, meaning that the sites obtained offer the best compromise between precision and recall. If the user wants to ensure that solutions will be found for highly pre-organized binding sites, they can run a low-tolerance calculation. This ensures high-precision results but obviously increases the risk of omitting relevant binding sites for which pre-organization is suboptimal. On the contrary, if the user works on non-optimized metal-binding sites (e.g., unbound X-ray structures or snapshots from MD simulations of apo proteins), calculations can be run in high-tolerance mode. In this case, more solutions are obtained, though more false positives may occur. However, we observed that this last option could explore transient sites and eventually map possible pathways for metal recruitment.

Internally, the program assigns a score threshold to each tolerance level. We parameterized these thresholds for each metal type by studying a random subset of structures from the pool used in the statistical analyses. For each structure, we asked BioMetAll 2.0 to identify all possible sites for the metal under study. For every score threshold between 0 and 1 (with an increment of 0.01), we compared BioMetAll’s results with the crystallographic metals. To consider a successful prediction, we used the same criterion than Dürr et al^11^: a distance between the predicted and crystallographic metal position below 5 Å. We counted true positives (sites appearing both in the prediction and the crystal), false positives (sites appearing in the prediction but not in the crystal), and false negatives (sites appearing in the crystal but not in the prediction). Finally, we assigned the score thresholds for low and medium tolerance levels based on the precision-recall curves (details are reported in the directory *04_tolerance_calibration* of the repository annexed), while the maximum tolerance is always assigned a score threshold of 0.

### Benchmark

Once we implemented the new scores and descriptors and properly parameterized the algorithm, we sought to assess BioMetAll 2.0’s efficiency in predicting crystallographic binding sites. To that end, we generated a test dataset comprising all structures deposited in the PDB^22^ between September 2019 and June 2023 that were not included in the statistical analysis. From them, we selected metal-binding sites with at least three coordinating amino acids, as these are the sites most likely to be biologically relevant, allowing us to compare our results with another state-of-the-art benchmark.^11^ The 2,871 crystallographic metal-binding sites finally included (list reported in the directory *05_validation* of the repository annexed) represent all the metals parameterized in the program except strontium.

For each case, we interrogated BioMetAll to identify all metal-binding sites for the specific metal present in the structure at a medium tolerance level. We ran the benchmark twice, varying the descriptors included (“AB” and “AB” + “FS”). We also considered the possibility of coordination with backbone oxygens. For each run, we counted true positives, false positives, and false negatives as in the parameterization step. Then, we calculated precision and recall for each metal and run (Table 3).

**Table 3.** Precision and recall values of the benchmark.

| Metal or group | Number of crystallographic binding sites | “AB” descriptors |  | “AB” + “FS” descriptors |  |
| --- | --- | --- | --- | --- | --- |
|  |  | Precision | Recall | Precision | Recall |
| Cd | 10 | 0.38 | 0.75 | 0.50 | 0.83 |
| Co | 21 | 0.73 | 0.91 | 0.89 | 0.89 |
| Cu | 108 | 0.89 | 0.83 | 0.87 | 0.72 |
| Fe | 279 | 0.38 | 0.35 | 0.69 | 0.66 |
| Hg | 1 | 1.00 | 1.00 | 1.00 | 1.00 |
| Mn | 230 | 0.54 | 0.56 | 0.88 | 0.84 |
| Ni | 45 | 0.54 | 0.43 | 0.92 | 0.80 |
| Zn | 908 | 0.85 | 0.91 | 0.93 | 0.90 |
| Ca | 712 | 0.54 | 0.56 | 0.92 | 0.83 |
| Mg | 257 | 0.32 | 0.39 | 0.76 | 0.79 |
| K | 66 | 0.28 | 0.24 | 0.77 | 0.25 |
| Na | 234 | 0.26 | 0.21 | 0.70 | 0.26 |
| Transition | 1602 | 0.66 | 0.81 | 0.84 | 0.92 |

The benchmark indicates that combining backbone and side-chain descriptors (“AB” + “FS”) provides high accuracy, with precision and recall exceeding 65%. Zinc predictions are especially precise, reaching about 90% in both recall and precision, which aligns with the larger number of structures in the training set. Overall, positive trends are also seen across all transition metals. Using only the backbone descriptor (“AB”) yields acceptable results for most metals, although it loses roughly 20% compared to the combined “AB” + “FS” approach. Some metals, particularly iron and nickel, are more affected, with iron showing a roughly 40% decrease in performance. This may be due to iron’s complex electronic behavior, including redox properties, initial coordination geometry, and spin states, which likely require case-by-case benchmarks beyond X-ray resolution. Based on this consistent behavior across transition metals, we created a “Transition” group as an alternative for certain applications, such as identifying iron-binding sites or screening large protein datasets. Predictions for both backbone and first coordination sphere are reliable in terms of precision and recall.

### Expanding towards non-transition metals

Expanding our benchmark to include alkaline and alkaline earth elements, we observed good results for alkaline earth metals (Ca and Mg). The predictions were particularly accurate when using the side-chain approximation, despite a roughly 50% decline in backbone-descriptor accuracy. This indicates that, generally, the backbone’s pre-organization is less influential for these metals than for transition metals. The most challenging cases involved alkaline elements, with potassium and sodium showing recall rates below 30%, although their side chain prediction precision remained above 65%. For these metals, alternative hypotheses should be explored to better understand and improve the low prediction quality.

## Conclusion

The backbone pre-organization that underpins the initial version of BioMetAll proved effective at detecting potential metal-binding sites, even when the geometries are not optimal. When experimental data on the metal’s exact position is absent, this approach seems especially valuable. In fact, this is a crucial advantage in designing new-to-nature metalloenzymes or metallopeptides, as it provides a more flexible scaffold that can be further refined through experimental methods such as directed evolution or other chemical or biochemical techniques.

However, backbone-based prediction has its limits; it does not account for more subtle effects that could contribute to the completeness of metal-protein interactions, which leads to uncertainty about the true site, transient sites, and noise. By providing a set of optimized scoring functions for BioMetAll in this new version and enabling both sidechain and backbone predictions, the risk of false positives is reduced, although the information obtained by both approaches could be particularly valuable. For example, calculations with a backbone and high tolerance allow the identification of putative sites on structures that do not show the full extent of pre-organization of the side chain resulting from metal binding.

## Acknowledgments

The authors are thankful for financial support from the Generalitat de Catalunya (grant 2021 SGR-866) and the Spanish Ministerio de Ciencia e Innovación (PID2023-149492NB-I00).

## Repository

Data associated with the manuscript, including a script for a visual interface in ChimeraX, will be available at https://github.com/insilichem/biometallv2.

Raúl Fernández DÍaz present address: BM Research, Dublin, Dublin D15 HN66, Ireland; School of Medicine, University College Dublin, Dublin D04 C1P1, Ireland; Conway Institute of Biomolecular and Biomedical Science, University College Dublin, Dublin D04 C1P1, Ireland; The Research Ireland Centre for Research Training in Genomics Data Science, Ireland

